# Evolutionary conserved and divergent responses to copper zinc superoxide dismutase inhibition in plants

**DOI:** 10.1101/2023.02.11.528141

**Authors:** Stephanie Frohn, Fabian B. Haas, Bernd H. Dreyer, Erik V. Reiss, Anne Ziplys, Heiko Weichert, Benjamin G. Chavez, John C. D’Auria, Stefan A. Rensing, Jos H.M. Schippers

**Author notes:** Corresponding author: Jos H.M. Schippers.

## Abstract

Life evolved in the presence of reactive oxygen species (ROS) and was further challenged by two consecutive great oxidation events. Therefore, ROS are deeply intertwined into the physological, morphological and transcriptional responses of organisms. Copper zinc superoxide dismutases (CuZnSODs) evolved around the first great oxidation event and have next to their classical role in ROS detoxification also important roles in signaling and transcriptional regulation. Here we addressed the role of CuZnSODs in early land plant evolution. We show, that pharmaceutical inhibition of CuZnSODs with Lung Cancer Screen 1 (LCS-1) in different plant species, including *Marchantia polymorpha* and *Physcomitrium patens*, representing the evolutionary early stages of land plants, and *Arabidopsis thaliana* as a modern vascular plant, lead to impairment of development and growth. Interestingly, Marchantia only possesses the cytosolic CuZnSOD isoform, whereas *Physcomitrium* additionally contains a plastidial isoform and Arabidopsis contains next to that a third peroxisomal isoform. An RNA-seq analysis revealed that the inhibition of CuZnSODs provoked a similar core response in all plant species analyzed, while those that contain more isoforms showed an extended response. In addition, an untargeted metabolomics approach revealed a specific metabolic signature for each plant species. Through the above approach the oxidative stress provoked by LCS-1 in plants can be specified and we argue that CuZnSOD functions are evolutionary conserved and might be important for plant terrestrialization.

## 1. Introduction

The identification of the enzyme superoxide dismutase (SOD) in red blood cells (McCord and Fridovich, 1969), represents a turning point in cell biology, leading to the field of reactive oxygen species (ROS) and redox biology as we know today (Dreyer and Schippers, 2019). SOD proteins convert superoxide (O_2_^•−^) to the relative stable and less toxic hydrogen peroxide (H_2_O_2_) and oxygen. Although O_2_^•−^ can spontaneously dismutate, cells need to scavenge it as uncontrolled formation will result in unwanted reactions with cellular components (e.g. metal ions and iron–sulfur clusters in proteins), resulting in the formation of highly damaging hydroxyl radicals (Wang et al., 2018). SODs are found in the vast majority of organisms that live in the presence of oxygen, as these enzymes represent the first line of defense against oxygen-derived free radicals (Dreyer and Schippers, 2019). Three types of highly conserved SOD enzymes evolved, that are classified by the catalytic metal ions they contain, resulting in MnSODs/FeSODs, NiSODs and Cu/ZnSODs (Wang et al., 2018). Amongst these, Cu/ZnSODs evolved relative late, during the time of the Great Oxidation Event (Inupakutika, et al., 2016; Dreyer and Schippers, 2019). The three independently evolved SOD types highlight the essential roles that these proteins play in different organisms. Moreover, nearly all streptophytes employ Fe- Mn- and CuZn-SODs, indicating the fundamental importance of these proteins for photosynthetic organisms (Dreyer and Schippers, 2019).

CuZnSOD enzymes act as homodimer in which each subunit binds a copper and zinc ion in the native state. However, heterologous expressed and purified CuZnSOD often binds a non-stoichiometric amount of the metal ions (Tajira et a., 2022), indicating that folding and metal ion loading of the enzyme is highly regulated *in vivo.* It has been shown that the redox state is a major regulator of this process as only the disulfide-reduced state of apo-SOD was monomeric and able to bind only one zinc ion per monomer (Tajira et a., 2022). Furthermore, intracellular CuZnSOD obtains its copper ions through the highly conserved COPPER CHAPERONE FOR SOD1 (CCS), which also introduces a disulfide bond that results in protein maturation (Culotta et al., 1998; Abdel-Ghany et al., 2005). Eukaryotic organisms contain up to 3, potentially 4 isoforms of CuZnSODs. Plants, yeast and animals all contain a soluble isoform that is mainly in the cytosol, and referred to as SOD1 in animals and CSD1 in plants. The second isoform, CSD2 is only found in plants (Dreyer and Schippers, 2019), and localizes to plastids. The third isoform, CSD3/SOD3, differs at sequence level the most from CSD1 and CSD2, and is found as an extracellular protein in yeast and animals, but as a peroxisomal protein in plants. In recent years, exciting discoveries where made concerning the cytosolic isoform, which in animals and yeast were found to act in the nucleus as a potential transcriptional regulator (Tsang et al., 2014; Li et al., 2019). Still, the exact role of each isoform is not completely understood.

In the field of medicine, SOD1 became of major interest as mutations in the protein were found to cause amyotrophic lateral sclerosis (ALS) (Rosen et al., 1993). 30 years later it is still heavily debated how the mutated forms of SOD1 cause ALS, and significant research activities regarding this remain ongoing. In addition, SOD1 is an important target for cancer therapy, as it maintains ROS homeostasis to support oncogene-dependent proliferation (Somwar et al., 2011; Gomez et al., 2019). In yeast, loss of SOD1 or CCS causes severe DNA damage and replication arrest under aerobic conditions (Carter et al., 2005). In plants, CuZnSODs have been mainly molecular characterized in the model species *Arabidopsis thaliana.* It has been shown that CSD genes are involved in the response to abiotic stress pathways and potentially also in developmental aspects of the plant (Dreyer and Schippers, 2019), still many open questions remain regarding their exact role in plants and how conserved their function within the streptophytic lineage is.

To obtain a better understanding of the role of CuZnSOD in the oxidative stress response pathways in plants, a comparative analysis of the effect inhibition of CuZnSOD activity in three plants species, *Arabidopsis thaliana*, *Marchantia polymorpha* and *Physcomitrium patens*, was analyzed. To this end we employed the small chemical lung cancer screen-1 (LCS-1), which was shown to primarily act as an inhibitor of CuZnSOD activity and to cause oxidative stress (Somwar et al., 2011). Our transcriptome and metabolome analysis reveal common and divergent responses amongst the three plant species tested. Which is especially of interest when considering that *Marchantia polymorpha* contains only one CuZnSOD isoform. Interestingly, we observe a strong transcriptional effect on glutathione biosynthesis and glutathione S-transferases, amongst all plants tested, indicating a link between the maintenance of ROS homeostasis and the activity of CuZnSOD. Based upon the results the essential roles of each CuZnSOD isoform are discussed and placed into an evolutionary context.

## 2. Material and methods

### 2.1. Plants and cultivation media

LCS-1 was dissolved in dimethyl sulfoxide (DMSO) to a stock concentration of 50mM and used for the treatments according to the final concentrations of 10μM or 50μM depending on the experiment. For phenotypic comparison LCS-1 was added to the solid medium. Mock treatment was performed with the same amount of DMSO for the same duration. The plants were grown depending on the species in different liquid or solid medium. Arabidopsis thaliana Col-0 seedlings were cultivated for 14 days in ½ MS + 1% sucrose (Murashige and Skoog, 1962), Physcomitrium patens type Reute for 21 days in Knop + microelements and Marchantia polymorpha type BoGa in ½ B5 + 1% sucrose for 21 days under long day condition (16/8) and at 21 degree at daytime, 19 degree at night time (Busch et al., 2019).

Plant material used for RNA-Seq and GCMS analysis was cultivated in liquid medium for the duration as described above without LCS-1 addition. For the treatment the medium was supplemented in the morning with LCS-1 dissolved in DMSO to a final concentration of 50μM or DMSO only for mock treatment. The cultivation of the plants was continued for 6 or 24 hours under the same conditions. Within the harvesting process the material was cleaned from medium residues and quick frozen.

### 2.2 Cloning, protein expression and purification

For the heterologous expression of AtCSD1 the CDS was cloned into the pET23a vector and transformed into the E. coli BL21 pLysS strain. The recombinant AtCSD1 protein containing a 6xHis-tag was purified using Dynabeads His-Tag (Thermo Fisher Scientific) according to manufacturer’s instructions. Five hundred microliters cleared extract was mixed with 500 μl binding buffer (50 mM NaP, pH 8.0, 300 mM NaCl, 0.01% Tween-20), and 50 μl washed Dynabeads were added. After 10 min incubation on a roller, the beads were washed 7 × with binding buffer, and 7 × with binding buffer, 5 mM imidazole. The elution was done with binding buffer, 150 mM imidazole, and the protein concentration was determined by performing a Bradford assay.

### 2.3 Superoxide dismutase activity assay

SOD activity was measured by a photometric approach adapted from Janknegt et al. (2007) using the color change by NBT-formazan formation in the absence of SOD activity. The basic reaction buffer (SOD buffer) was adapted to 100μl 0.1 M potassium phosphate buffer (KH2PO4), 0.4 μl 65 mM methionine, 4 μl 3.75 mM NBT and 67.06 μl ddH2O. The SOD sample volume was adjusted to 20 μl and in addition with SOD buffer and 0.5 mM riboflavin dissolved in 0.1 M potassium phosphate buffer incubated for 10 s in the dark before illumination for 15 min with shaking on a fluorescence light source. Chemical blanks consisting of 175 μl SOD buffer, 5 μl riboflavin and 20 μl KH2PO4 buffer as well as sample blanks consisting of 190 μl KH2PO4 buffer and 20 μl sample are treated in the same way. Finally, the absorbance was measured at 560 nm in a 6×6 full circle (1500 μm border) and with 30 s shaking and 30 s waiting between different timepoints. The SOD activity was calculated as percentage inhibition of NBT-formazan formation in comparison to chemical blank.

### 2.4 RNA extraction and RNA-sequencing

RNA extraction of Marchantia polymorpha and Physcomitrium patens was done using the Spectrum^™^ Plant Total RNA Kit (Sigma) and for Arabidopsis thaliana the NucleoSpin RNA Plant min kit (Macherey -Nagel) was used. RNA was extracted according to the manufacturer’s recommendations. Libraries were generated using DNA-free RNA with the NEBNext^®^ Ultra^™^ II RNA Library Prep Kit for Illumina according to manufacturer’s instructions. Sequencing was performed on an Illumina HiSeq2000 in 150-bp pair-end mode at Novogene (London, UK) with a minimum of 30 million reads per biological replicate.

### 2.5 RNA-seq data analysis

RNA-seq data analysis was largely performed as previously described (Pierre-Francois Perroud et al., 2018). Read mapping was done using GSNAP (Wu and Nacu, 2010), read count was done using HTSEQ-COUNT against the respective organism gene models (Anders et al., 2015). Differential Expressed Genes (DEGs) were called using three different algorithms, namely EGDER (Robinson et al., 2010), DESEQ2 (Love et al., 2014) and NOISEQ (Tarazona et al., 2011). Subsequently the overlap genes between the 3 different algorithms were taken as DEGs and used in subsequent downstream analysis.

### 2.5 Metabolite extraction and GC-MS analysis

Polar metabolites were extracted from deep-frozen homogenized plant material using a previously described extraction protocol (Lisec et al., 2006). The protocol was implemented on a liquid handling system (Biomek^®^ FXP, Beckman Coul-ter GmbH, Krefeld, Germany): 0.625 ml chilled extraction buffer (2.5:1:1 v/v MeOH/CHCl_3_/H_2_O) and 0.250 ml H_2_O after incubation. Extracts from eight biological replicates were processed into aliquots containing 50 μl of polar phase sample. Dried extracts were in-line derivatized prior to injection using a Gerstel MPS2-XL autosampler (Gerstel, Germany) and analyzed in split mode (1:3) using a LECO Pegasus HT time-of-flight mass spectrometer (LECO, USA) hyphenated with an Agilent 7890 gas chromatograph (Agilent, USA). Sample identification of known and unknown features occurred using the LECO ChromaTOF software package while using the Golm Metabolome Database (GMD; Kopka et al., 2005). Peak intensities were determined using the R package TargetSearch and normalized regarding to sample weight and measuring day/detector response (Cuadros-Inostroza et al., 2009).

## 3. Results

### 3.1 Phylogenetic relationship between CuZnSODs

As noted previously, CuZnSODs are highly conserved enzymes amongst eukaryotes (Dreyer and Schippers, 2019). LCS-1 has so far been shown to be effective in inhibiting SOD1 from humans and mice (Somwar et al., 2011; Li et al., 2018), and the high homology with plant CSD proteins suggest that it might also be an effective compound in plants (Fig. 1). In *Arabidopsis thaliana* and *Oryza sativa*, three CuZnSODs isoforms can be found, namely the cytosolic CSD1, the plastidial CSD2 and the peroxisomal CSD3. Interestingly, the here selected gymnosperms, *Ginkgo biloba* and *Pinus sylvestris*, only have the cytosolic and plastidial isoforms of CSD. Furthermore, the *hornwort Anthoceros punctatus* and the moss *Physcomitrium patens* also contain only the cytosolic and plastidial isoforms, but in the later for each isoform two copies are present. In contrast, the freshwater algea *Chara braunni*, the liverwort *Marchantia polymorpha* and the fern *Azolla filiculoides* all contain only the cytosolic CSD isoform. These differences in the number of CSD isoforms of plants is exploited here, by determining the impact of LCS-1 on *Arabidopsis thaliana*, *Physcomitrium patterns* and *Marchantia polymorpha*.

**Fig. 1.**
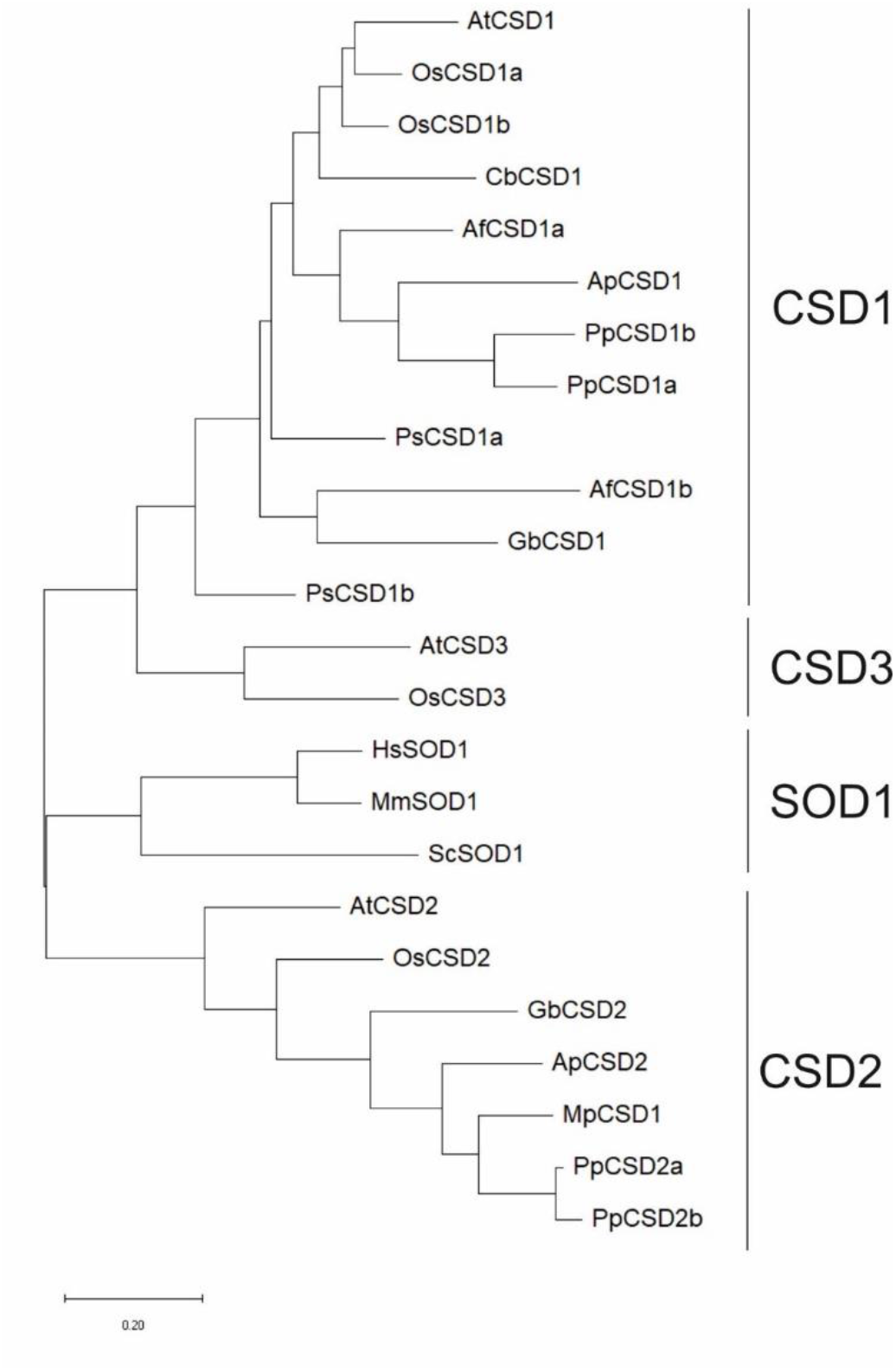
Phylogenetic relationship between CSD proteins in several selected species.

### 3.2 Purified CuZnSODs of plants are inhibited by LCS-1

In order to understand if LCS-1 is capable of inhibiting CuZnSOD activity of plant isoforms, AtCSD1 and MpCSD1 were heterologous expressed, purified and tested for their activity in an *in vitro* enzyme activity assay (Fig. 2A). To this end, AtCSD1 was expressed in *E. coli*, while MpCSD1 was expressed in in the yeast *Pichia pastoris*. Both proteins were purified and quantified using a Bradford assay. To determine the activity of both CSD proteins, we initially employed a standard activity assay that relies on xanthine oxidase to form the substrate for CSD. However, we observed like others that LCS-1 interferes with this assay (Malik et al., 2020), and therefore switched to an assay that relies on riboflavin for the formation of superoxide (Janknegt et al., 2007). Here we found that in the presence of 50 μM LCS-1, both MpCSD1 and AtCSD1 dramatically loose activity (Fig. 2.).

**Fig. 2.**
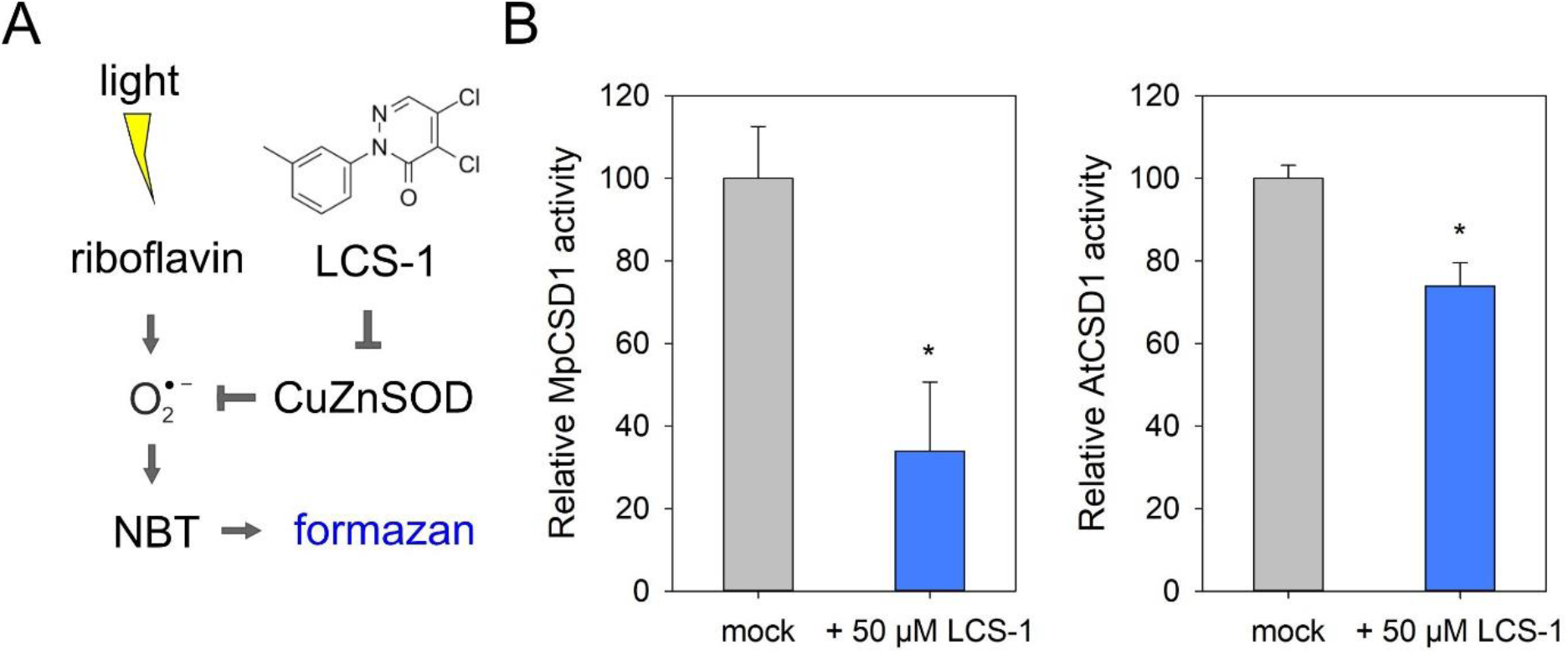
Activity of CSD proteins is inhibited by LCS-1 *in vitro.* (A) Principle of the RF-NBT assay to measure SOD activity in the presence of LCS-1. (B). MpCSD1 and AtCSD1 enzymatic activity was assayed *in vitro* under mock conditions or in the presence of 50 μM LCS-1. Relative activity of both plant SODs decreased in the presence of LCS-1. Bars indicate the mean ± SD of at least three measurements. Asterisk indicates significant difference as compared to the mock conditions according to student’s T-Test (P < 0.05).

### 3.3 LCS-1 inhibits plant growth in a concentration dependent manner

Loss of SOD1 activity in cancer cells results in growth inhibition and cell death (Somwaret al., 2011; Gomez et al., 2019). To determine if LCS-1 has an impact on plant growth and development, plants were cultured *in vitro* and transplanted on medium containing either 10 or 50 μM of LCS-1. After two weeks the impact on plant growth was visually analyzed and the dry weight was determined. Both male and female *Marchantia polymorpha* plants showed a strong growth suppression on medium containing LCS-1 (Fig. 3A). Which was also evident from the biomass accumulation upon stress treatment (Fig. 3B). Similarly, *Physcomitrium patens* and *Arabidopsis thaliana* plants showed a strong growth inhibition upon treatment with LCS-1 (Fig. 3C-F). Thus, also in plants LCS-1 results in a severe growth arrest.

**Fig. 3.**
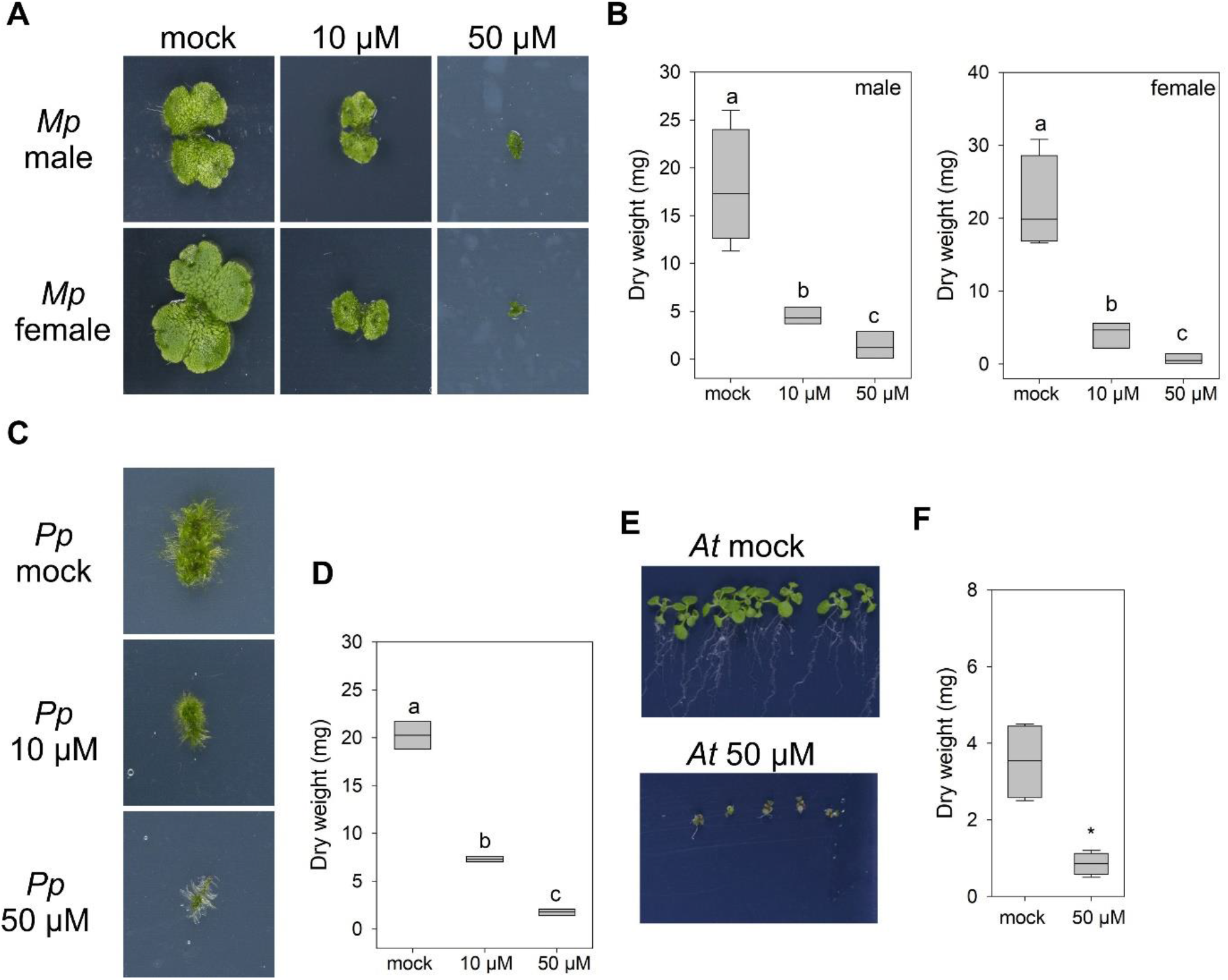
LCS-1 causes growth arrest in plants. (A) Images of *Marchantia polymorpha* plants grown for two weeks after transfer on either control medium (mock) or medium containing LCS-1. (B) The dry weight of *Marchantia polymorpha* plants exposed to different concentrations of LCS-1. Box plots indicates the average dry weight. Different letters above the boxes indicate significant difference to the controls (one-way ANOVA, p < 0.05). (C) *Physcomitrium patens* plants were imaged after transfer and 2-weeks of growth control medium (mock) or medium containing LCS-1. (D) Box plot indicates the average dry weight. Different letters above the boxes indicate significant difference to the controls (one-way ANOVA, p < 0.05). (E) Representative images of transferred *Arabidopsis thaliana* plants after 2 weeks of growth on control medium (mock) or LCS-1 containing medium. (F) Box plot indicates the average dry weight of Arabidopsis thaliana seedlings (n = 10). Asterisk indicates significant difference according to student’s T-Test (p < 0.05).

### 3.4 Transcriptional response to LCS-1 treatment reveals a strong induction of oxidative stress

At the physiological level, all plants show a similar growth arrest phenotype. To understand if at the molecular level a common response is initiated, a RNA-seq based transcriptome profiling was performed after 6 and 24 hours of LCS-1 application. As the number of CSD isoforms differ between species, we expected to observe common and divergent responses between the selected species.

Analysis of the transcriptome data for *Marchantia polymorpha* resulted in the identification of 152 DEGs in male and/or female plants at both stress timepoints (Fig. 4A; Table S1). After 6 hours of LCS-1 treatment, 70 DEGs in male plants were detected while in female plants 98 DEGs were detected. Of the 70 DEGs in male plants, 65 were significantly induced, while 5 were downregulated. In female plants, 61 genes were upregulated and 37 genes were downregulated. In general, genes in female plants tended to respond stronger to 6 h of LCS-1 treatment as compared to male plants (Fig. 4A). After 24 hours, most upregulated genes are decrease their expression level in male and female plants. In male plants, 64 DEGs are detected of which 43 are upregulated and 21 are downregulated. In female plants, 23 DEGs were still present after 24 h of LCS-1 of which 21 are upregulated and 3 downregulated. A gene ontology (GO) enrichment analysis of the DEGs mainly indicated that glutathione metabolic processes and glutathione transferase activity are among the few terms that are significantly and strongly enriched (Fig. 4B). A close examination of this group revealed that the glutathione biosynthesis gene *MpGSH1* and the *GLUTATHIONE REDUCTASE 1* gene *MpGR1*, are both induced during the initial phase of stress. Furthermore, 10 glutathione transferase (GST) genes are strongly induced upon LCS-1 treatment. As glutathione is essential for oxidative stress tolerance and the regulation of developmental processes (Jozefczak et al., 2012; Diaz-Vivancos et al., 2015), it indicates that LCS-1 provokes oxidative stress in *Marchantia polymorpha.* As CuZnSODs are needed to prevent superoxide-dependent lipid peroxidation (del Río et al., 2018), it is of interest to note that two genes involved in the final step of fatty acid biosynthesis are upregulated (Mp5g15340 and Mp3g12040), together with a gene encoding a lipid catabolic enzyme *NON-SPECIFIC PHOSPHOLIPASE C1* (Mp8g11280). These genes suggest a higher turnover of lipids upon LCS-1 treatment.

**Fig. 4.**
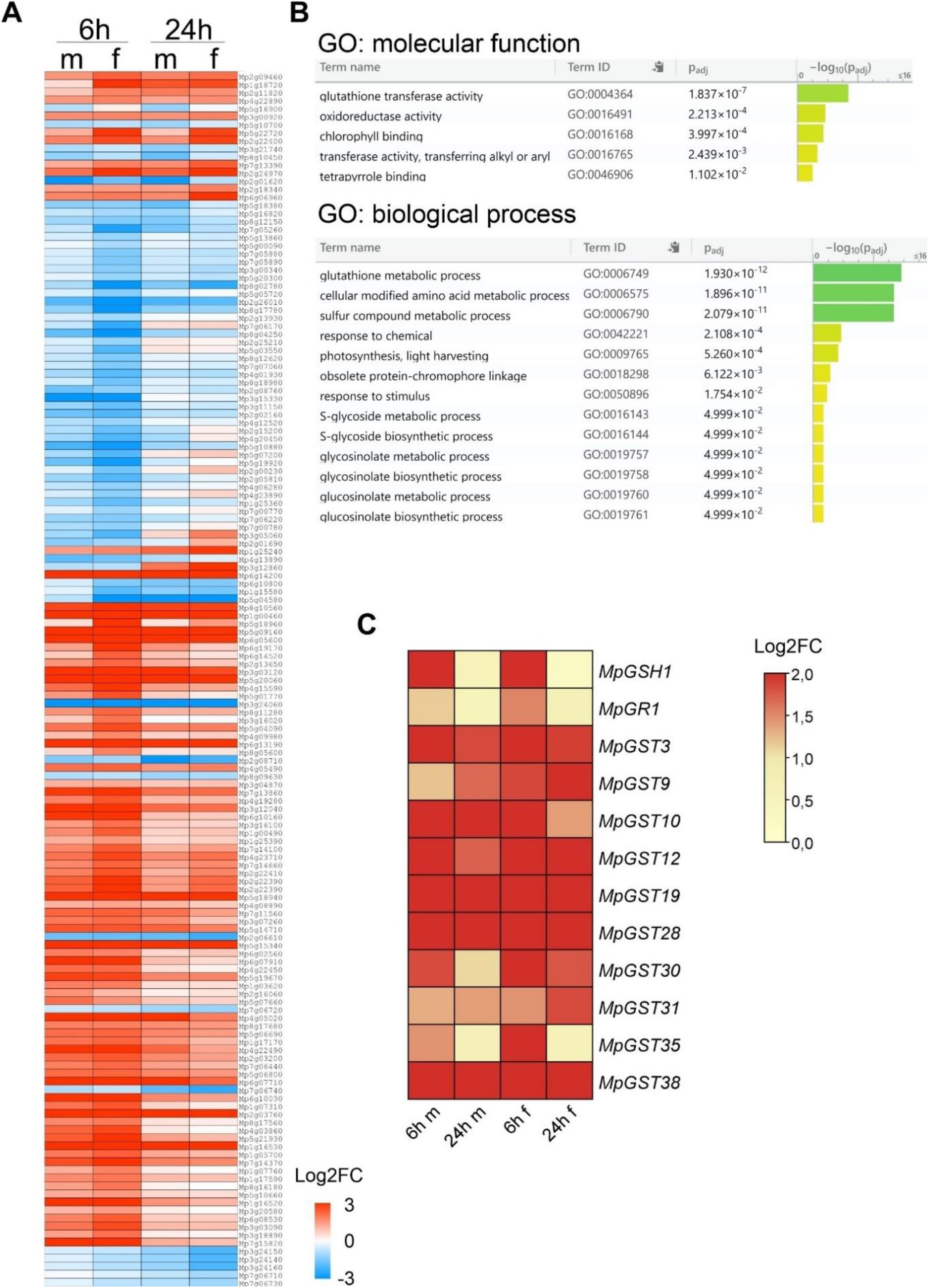
Transcriptional response of *Marchantia polymorpha* towards LCS-1. (A) Heatmap representation of the 152 DEGs significantly responding to LCS-1 in male and female plants. (B) A GO enrichment was performed using the g:Profiler tool (Raudvere et al., 2019), based on the ENSEMBL gene annotation for all 152 identified DEGs. (C) The expression of genes involved in glutathione biosynthesis and metabolism upon LCS-1 are depicted in a heatmap.

The response of the 152 LCS-1 DEGs was compared to published Marchantia transcriptome data under abiotic stress (Tan et al., 2021). This analysis revealed that most genes responding to LCS-1 stress are also responding to cold, high light and nitrogen deficiency (Table S1). This suggest that a large part of the genes are general stress genes, which might have in common to respond to alterations in ROS homeostasis under these conditions.

LCS-1 treatment of Physcomitrium resulted in the upregulation of 693 genes after 6h of stress and of 302 genes after 24 hours of stress, of which 155 overlap between both timepoints (Fig. 5A; Table S2). A GO enrichment analysis revealed, like for Marchantia, glutathione transferase activity and glutathione metabolic process to be significantly enriched (Fig. 5A). In addition, molecular function terms related to protein folding and phenylpropanoid metabolism are enriched. The biological process signature clearly indicates that the plants experience stress, as several abiotic stress response terms are enriched, including those for oxidative stress, response to reactive oxygen species and response to hydrogen peroxide. Furthermore, genes involved in organic acid metabolic processes and secondary metabolism are enriched upon LCS-1 treatment. Next to that, 1035 genes are downregulated after 6h of LCS-1 treatment, and 860 after 24h of stress treatment (Fig. 5B), of which 462 genes overlap. Like Marchantia, genes related to chlorophyll binding and tetrapyrrole binding are enriched among the downregulated genes. Indicating that the LCS-1 stress impairs photosynthetic processes. This might be especially relevant for *Physcomitrium patens* as it contains two plastidial CSD isoforms (Fig. 1). Interestingly, genes related to cell wall biosynthesis and organization are downregulated upon LCS-1 treatment. The altered expression of cell wall related genes might indicate that apoplastic ROS homeostasis is disturbed upon LCS-1 treatment, which is required for the regulation of cell growth and deposition of secondary cell walls (Schmidt et al., 2016).

**Fig. 5.**
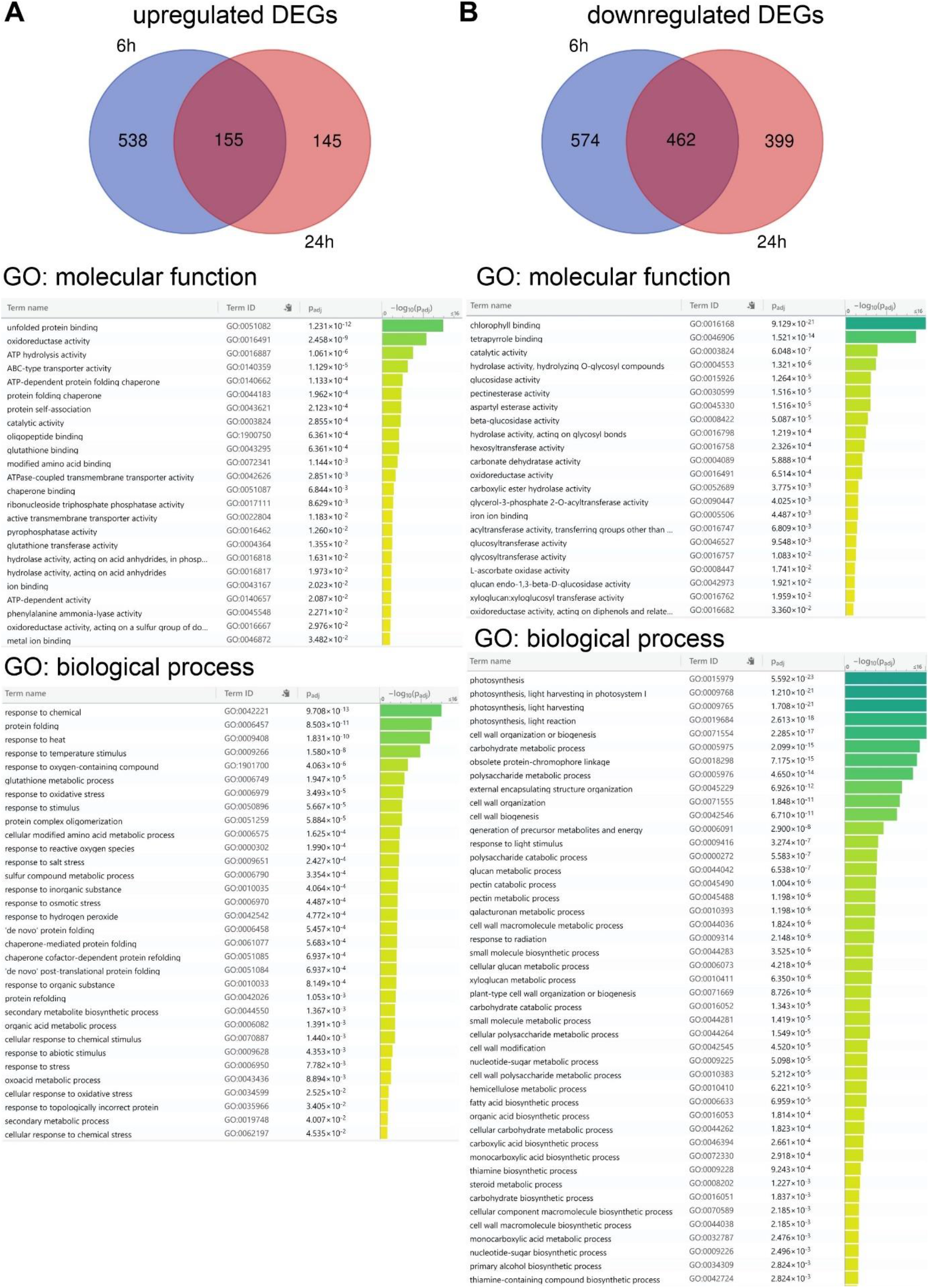
Transcriptional response of *Physcomitrium patens* towards LCS-1. (A, B) Venn diagrams depict the number of up- or downregulated DEGs after 6h and 24h of LCS-1 treatment and their overlap. In addition a GO enrichment analysis was performed using the g:Profiler tool (Raudvere et al., 2019), based on the ENSEMBL gene annotation for all DEGs. Shown are the enriched molecular function and biological process terms amongst the up- or downregulated genes.

The exposure of Arabidopsis towards LCS-1 resulted in the up-regulation of 148 genes after 6h and 672 genes after 24h, respectively (Fig. 6A; Table S3). Amongst these, 116 genes overlap, indicating that many of the initially induced genes remain highly expressed. As observed in *Marchantia polymorpha* and *Physcomitrium patens*, also in Arabidopsis genes belonging to glutathione transferase activity and glutathione metabolic process are strongly enriched upon LCS-1 treatment. However, no terms related to protein folding stress are enriched in Arabidopsis but genes involved in vitamin B6 (pyridoxal 5’-phosphate). Vitamin B6 is an excellent antioxidant against the superoxide anion (Jain and Lim, 2001). Furthermore, ABC transporters are activated, which are known to be able to transport herbicides in plants (Xi et al., 2012). The GO terms for biological process clearly indicate that the plants experience severe stress. Next to oxidative stress, especially terms related to low oxygen stress and hypoxia are enriched, which might indicate that energy metabolism is substantially impaired upon LCS-1 treatment. Among the upregulated genes are *PYRUVATE DECARBOXYLASE 1* (*PDC1*), *HUP40*, *HUP44*, *ANAC102*, *HYPOXIA RESPONSIVE ERF 2* (*HRE2*), *LOB DOMAIN-CONTAINING PROTEIN 41* (*LBD41*), and *ALCOHOL DEHYDROGENASE 1* (*ADH1*), which are typical genes to be induced upon low oxygen stress and an energy crisis (Schmidt et al., 2018). Taken together, LCS-1 provokes a severe stress on plants that is potentially initiated by the inhibition of CuZnSODs, followed by oxidative stress and a potential collapse of energy metabolism. After 6h of LCS-1 treatment 68 genes are significantly down regulated (FC >−2; p < 0.05), while after 24h, 183 genes are downregulated (Fig. 6B). Amongst both time points only 15 genes overlap. Both photosynthetic processes as well as tetrapyrrole binding are also downregulated in Arabidopsis, as observed for *Marchantia polymorpha* and *Physcomitrium patens*. Furthermore, secondary metabolism processes are impaired, including glucosinolate biosynthesis and coumarin biosynthesis. Taken together, the three analyzed plant species show common and differential responses to LCS-1. The responses observed in Marchantia are largely conserved in *Physcomitrium* and Arabidopsis, but the latter two species show additional responses that might be related to the additional CuZnSOD isoforms they contain.

**Fig. 6.**
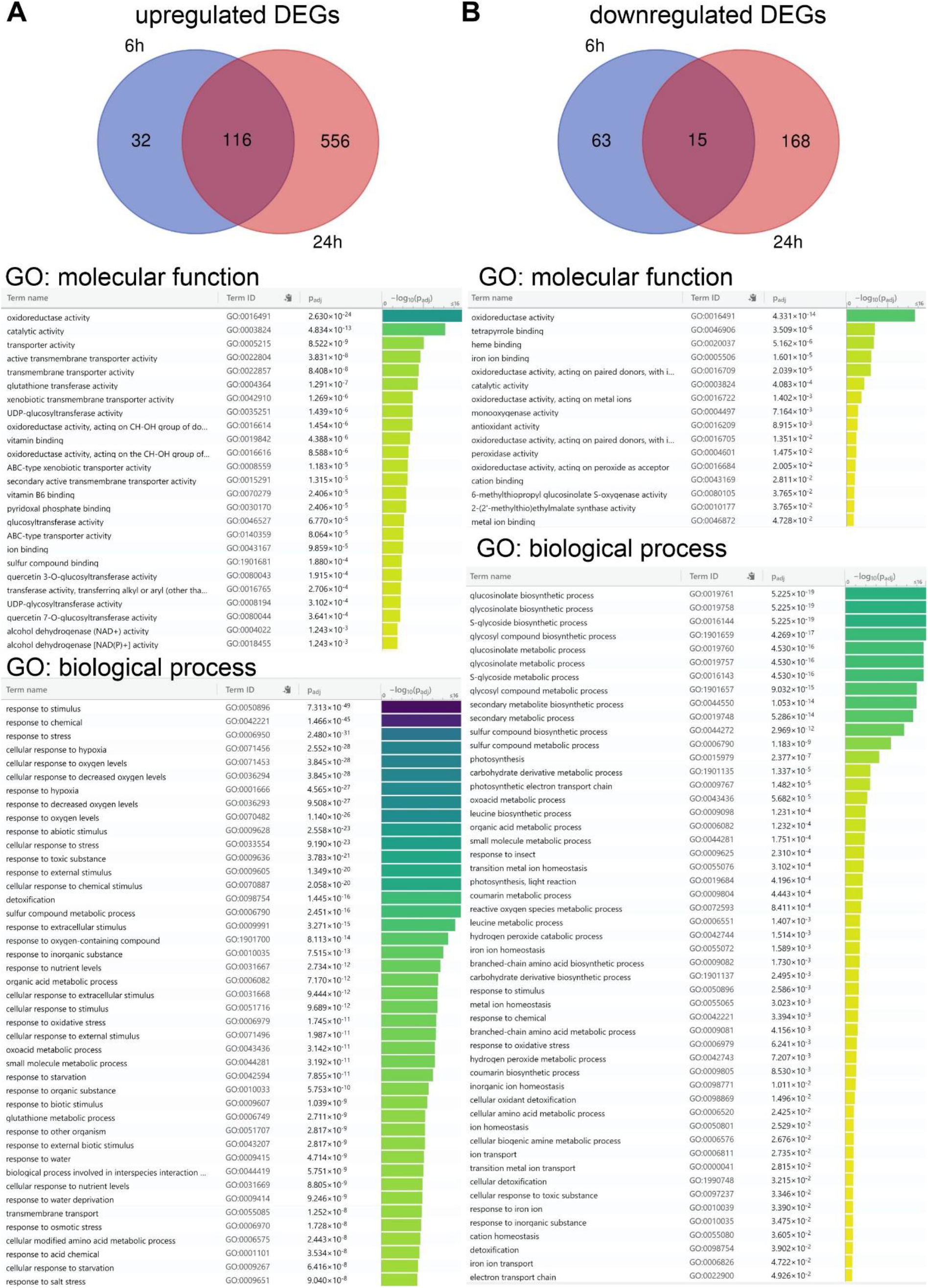
Transcriptional response of *Arabidopsis thalian* to LCS-1 stress. (A, B) Venn diagrams depict the number of up- or downregulated DEGs after 6h and 24h of LCS-1 application and their overlap. Below the results of a GO enrichment analysis using the g:Profiler tool (Raudvere et al., 2019) are shown, based on the TAIR gene annotation for all DEGs. Shown are the enriched molecular function and biological process terms amongst the up- or downregulated genes.

### 3.5 LCS-1 affects different metabolic processes in plants

As indicated by the transcriptome analysis, organic acid metabolism appears to be downregulated in *Physcomitrium* and Arabidopsis. Here we used a GC-MS based approach to identify and quantify the metabolic changes upon LCS-1. A principal component analysis (PCA) of the measured metabolites revealed that the metabolic differences between the species used are larger as the effect of LCS-1 on the metabolic profile (Fig. 7). Nonetheless, Several metabolites showed differential responses towards LCS-1, especially also depending on the plant species treated. In Arabidopsis, the steady-state level of TCA cycle intermediates is hardly affected by the LCS-1 treatment, except for oxaloacete which can give rise to the oxaloacetate/aspartate family of amino acids. Indeed, at the amino acid level an increase in isoleucine, lysine, and methionine is observed, which might relate to the altered oxaloacetate levels. In contrast, Marchantia and Physcomitrium showed a more severe alteration in TCA cycle Intermediates as Arabidopsis. Of note, the male and female Marchantia plants showed for several intermediates contrasting levels. At the amino acids level, *Physcomitrium* shows a characteristic upregulation of phenylalanine and aspartic acid after 24h of LCS-1 treatment. In Marchantia plants also changes in proline and glutamine levels are observed. Next to that, all treated plants accumulate the polyamine putrescine, which is a well-known stress-related compound. Furthermore, at the level of measured sugars it was noted that mainly Arabidopsis accumulates sugars with antioxidant activity like galactinol and raffinose. In addition, all plants showed an upregulation of talose. Tagatose levels only increased in the mosses and liverworts tested here. Taken together, the species tested show differential responses at metabolic level.

**Fig. 7.**
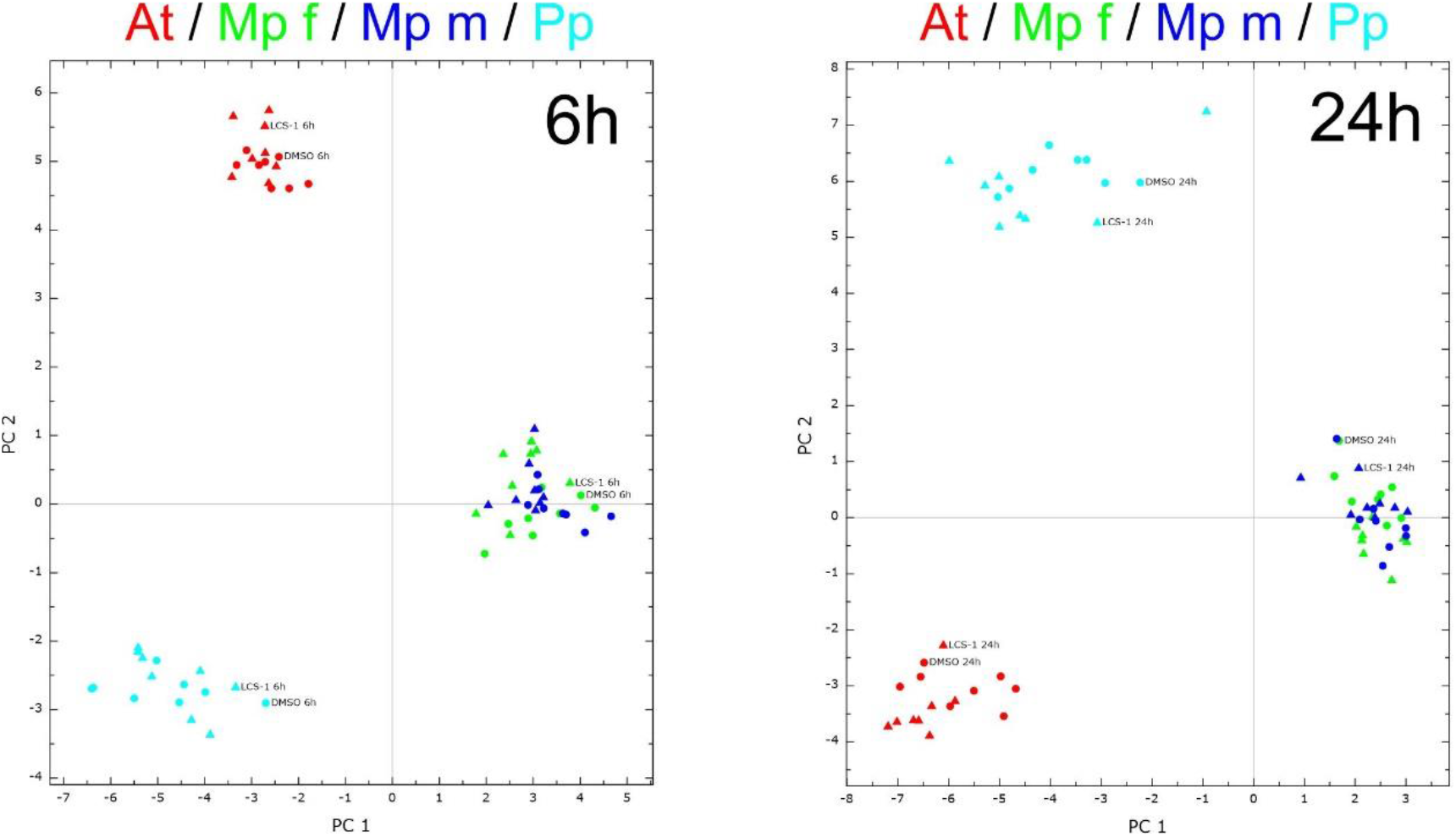
PCA analysis of the measured metabolism of the different plants mock treated or treated with LCS-1.

**Fig. 8.**
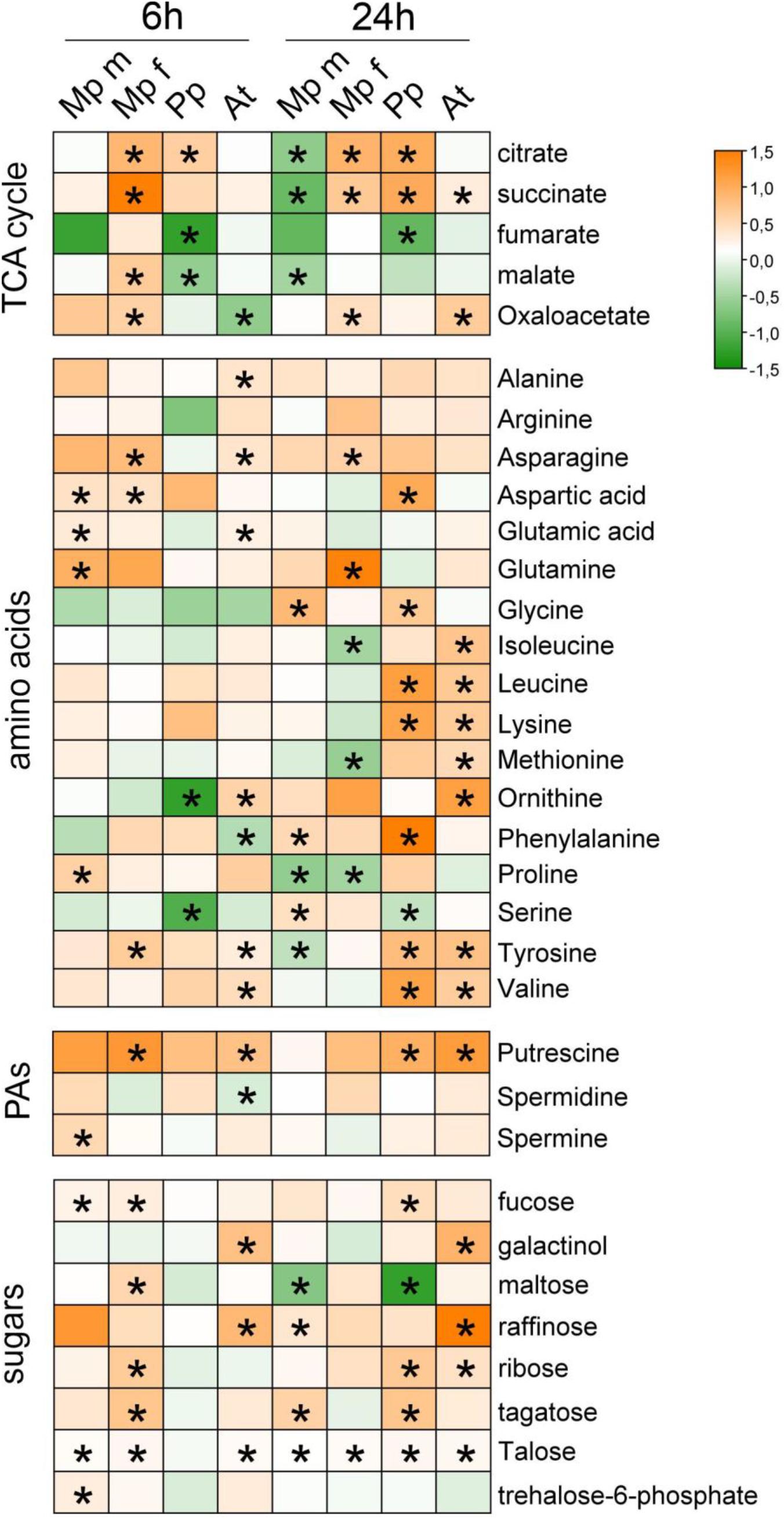
Metabolite profiles of the different plants is presented as a heat map. The colors indicate log2 fold change for the given metabolite. Asterisk indicates significant difference from mock (DMSO) treated plants (p < 0.05). Data shown is based on the measurement of 8 biological replicates.

## 4. Discussion

In this study, our transcriptomic and metabolic analyses demonstrated that the SOD inhibitor LCS-1 provokes oxidative stress in 3 different plant species. We show that LCS-1 is able to inhibit the enzymatic activity of plant CSD proteins, suggesting that the observed transcriptional responses are due to the inhibition of CuZn superoxide dismutase activity in plants. Like observed for animal cells, LCS-1 causes a severe growth arrest in plants.

CuZnSODs function as safeguards against a build-up of superoxide in cells (Dreyer and Schippers, 2019), which is able to readily react with metal ions, resulting in the formation of the hydroxyl radical, a major inducer of oxidative stress. As CuZnSOD proteins arose during the GOE-, especially in eukaryotic cells, it is likely that they were essential to the adaptation in an oxygenic environment. Especially, the water-to-land transition of plants resulted in a higher availability of oxygen, due to the more rapid diffusion in air. It is thought that land plants arose from freshwater algae, which all contain at least the cytosolic CSD isoform. Here, three land plant species were analyzed as they differ in the number of CSD isoforms and their position within the phylogenetic tree. Marchantia only contains a cytosolic CSD isoform, while Physcomitrium contains 2 cytosolic and 2 plastidial CSD isoforms, and Arabidopsis has 1 cytosolic, 1 plastidial and 1 peroxisomal isoform. To obtain insights into the role of the cytosolic CSD isoform and the expansion of CSD protein in plants, we used a chemical inhibitor against SOD, that was previously shown to be highly active against SOD1 from humans (Somwar et al., 2011).

The here performed biochemical and physiological analysis demonstrates that LCS-1 is also able to inhibit plant CuZnSOD isoforms. In the assay tested both MpCSD1 and AtCSD1 were shown to be sensitive to the addition of LCS-1 (Fig. 2). Furthermore, plant growth of Marchantia, Physcomitrium and Arabidopsis is nearly completely impaired on medium containing 10-50 μM of LCS-1 (Fig. 3). This inhibition of cell growth was also observed for tumor cells (Somwar et al., 2011; Ling et al., 2022), whereby cell death is triggered by higher concentrations (20 μM). Thus, LCS-1 is able to inhibit cell growth in both eukaryotic systems, humans and plants.

Our transcriptome analysis uncovers a common response between the liverwort Marchantia, the moss Physcomitrium and the angiosperm Arabidopsis. As only the cytosolic CSD isoform is present in all species tested, it suggests that the common transcriptional response is caused by the inhibition of the cytosolic CSD isoform. LCS-1 treatment results in a rapid upregulation of genes encoding for glutathione transferases and enzymes involved in the biosynthesis of glutathione. Superoxide can readily react with metal ions, thereby causing the formation of hydroxyl radical (Fenton reaction). This metal-induced cellular toxicity is counteracted by plants through the usage of glutathione (Jozefczak et al., 2012). Glutathione is not only able to reduce oxidized cellular component, but can also scavenge metals through its thiol group, or promote metal chelating as a precursor for phytochelatins (PCs). Therefore, it appears that cytosolic CuZnSOD is required to remove superoxide in the cytoplasm to prevent unwanted reactions with metal ions.

Both Arabidopsis and Physcomitrium display a strong downregulation of genes involved in photosynthesis (Fig. 5, 6). In Marchantia, only a few genes related to photosynthesis are affected. The strong and broad response in the other two species might be directly related to the presence of CSD in their plastids. This inhibition of plastidial CuZnSOD might be responsible for the observed transcriptional responses. Previously, it was shown that knock-down of *AtCSD2* results in a severe impairment of tolerance against high light stress (Xing et al., 2013). Moreover, a complete knock-out of AtCSD2 is potentially lethal (Rizhsky et al., 2003; Xing et al., 2013), which might explain the massive transcriptional response in Physcomitrium and Arabidopsis as compared to Marchantia upon LCS-1 treatment. Still, Physcomitrium also displays unique responses, which involve the response to protein folding stress and an upregulation of chaperones. Abiotic stresses, which have in common oxidative stress, are well known to cause protein dysfunction (Wang et al., 2004). Especially Heat-shock proteins and chaperones are responsible for maintaining protein folding, assembly, translocation and degradation. As the other plants do not display an upregulation of chaperone activity, it might be that the oxidative stress that LCS-1 causes in plants, is more severe in Physcomitrium as compared to the two other species. Although Arabidopsis is the only tested plant here that also contains a peroxisomal CSD isoform, it is not clear from the trancriptomic data if peroxisomal function is impaired. It might be that on the one hand the antioxidant capacity of the peroxisome is sufficient to withstand the loss of CSD3 activity. On the other hand, and potentially more likely, LCS-1 might not affect CSD3 as it is distinct from CSD1 and CSD2. It would be of interest to test the impact of LCS-1 on CSD3 activity in a follow-up study.

Furthermore, a metabolite profiling was performed on the three different plant species after LCS-1 treatment. This analysis revealed that Marchantia and Physcomitrium showed a more disturbed metabolite profile as compared to Arabidopsis. Moreover, Arabidopsis increased the levels of galactinol and raffinose, two sugars with antioxidant capacity (Song et al., 2016), while the others did not. In addition, all plant species appear to upregulate the polyamine putrescine, which previously was shown to improve oxidative stress tolerance in Arabidopsis (Alcázar et al., 2010). All in all, the metabolic profile of Arabidopsis is reminiscent of a plant experiencing oxidative stress. How Marchantia and Physcomitrium adjust their metabolism to oxidative stress is to date not well studied.

Thusfar, CuZnSOD inhibitors have not been applied to plants to study the role of CSD proteins. Here we demonstrate that LCS-1 is a useful chemical to study CSD protein function and identify molecular pathways that are linked to enzyme activity. We uncovered that maintaining glutathione homeostasis is a conserved biological mechanism to deal with impaired CuZnSOD activity in plants. We expect that the here provided datasets will be help to uncover further biological mechanisms linked to CuZnSOD activity in plants.

## Acknowledgments

The work was carried out in the framework of MAdLand (https://madland.science, DFG priority programme 2237), the authors are grateful for funding by the DFG (SCHI 1130/9-1 and RE1597/19-1).

## Appendix A. Supplementary data

Supplementary data to this article can be found online

